## Supporting Information S1 Figure for "Predicting precision grip grasp locations on three-dimensional objects"

Department of Experimental Psychology, Justus Liebig University Giessen, Otto-Behaghel-Str.10F, Giessen 35394, Germany

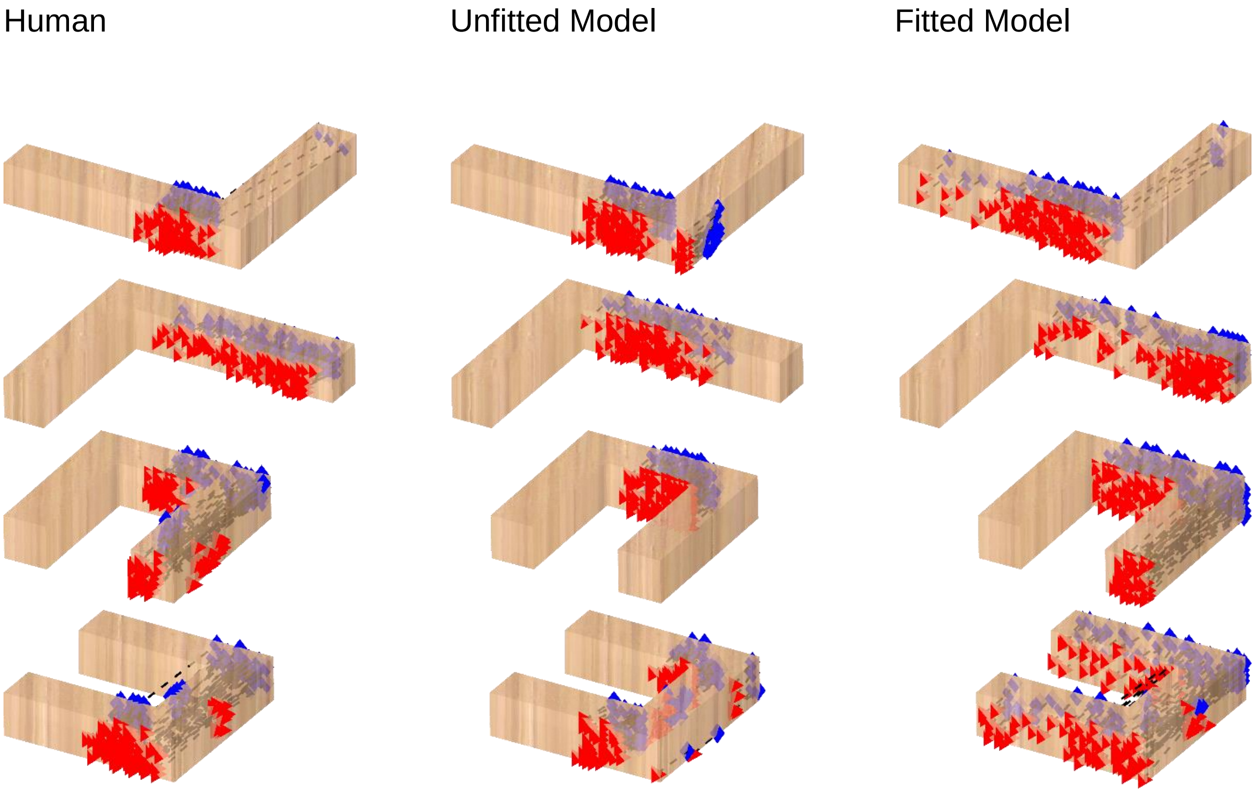

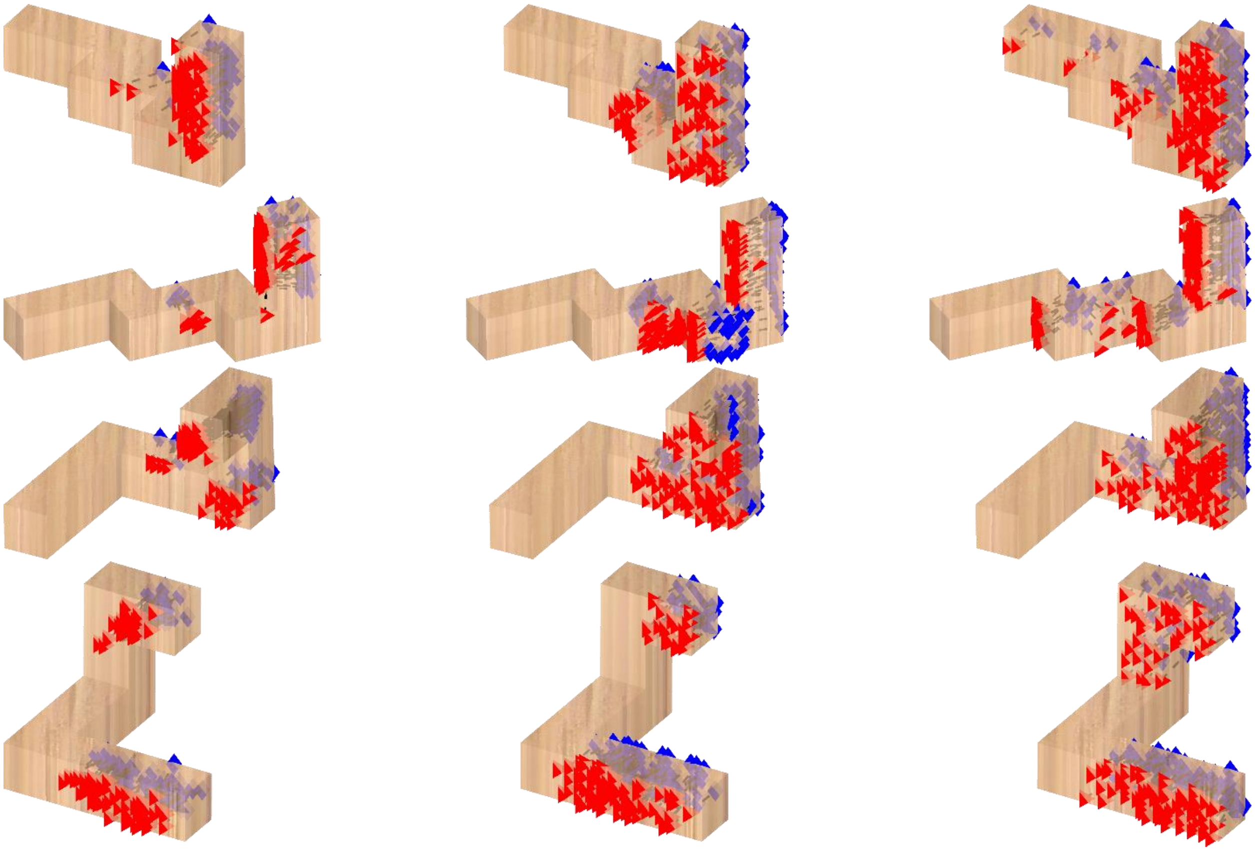

(b) Experiment 2

Human | Unfitted Model | Fitted Model

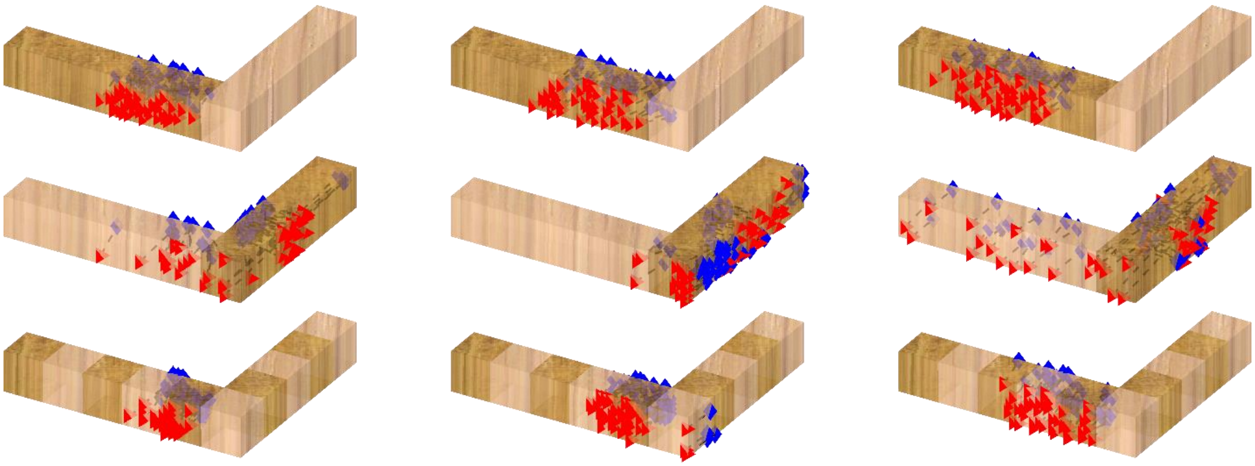

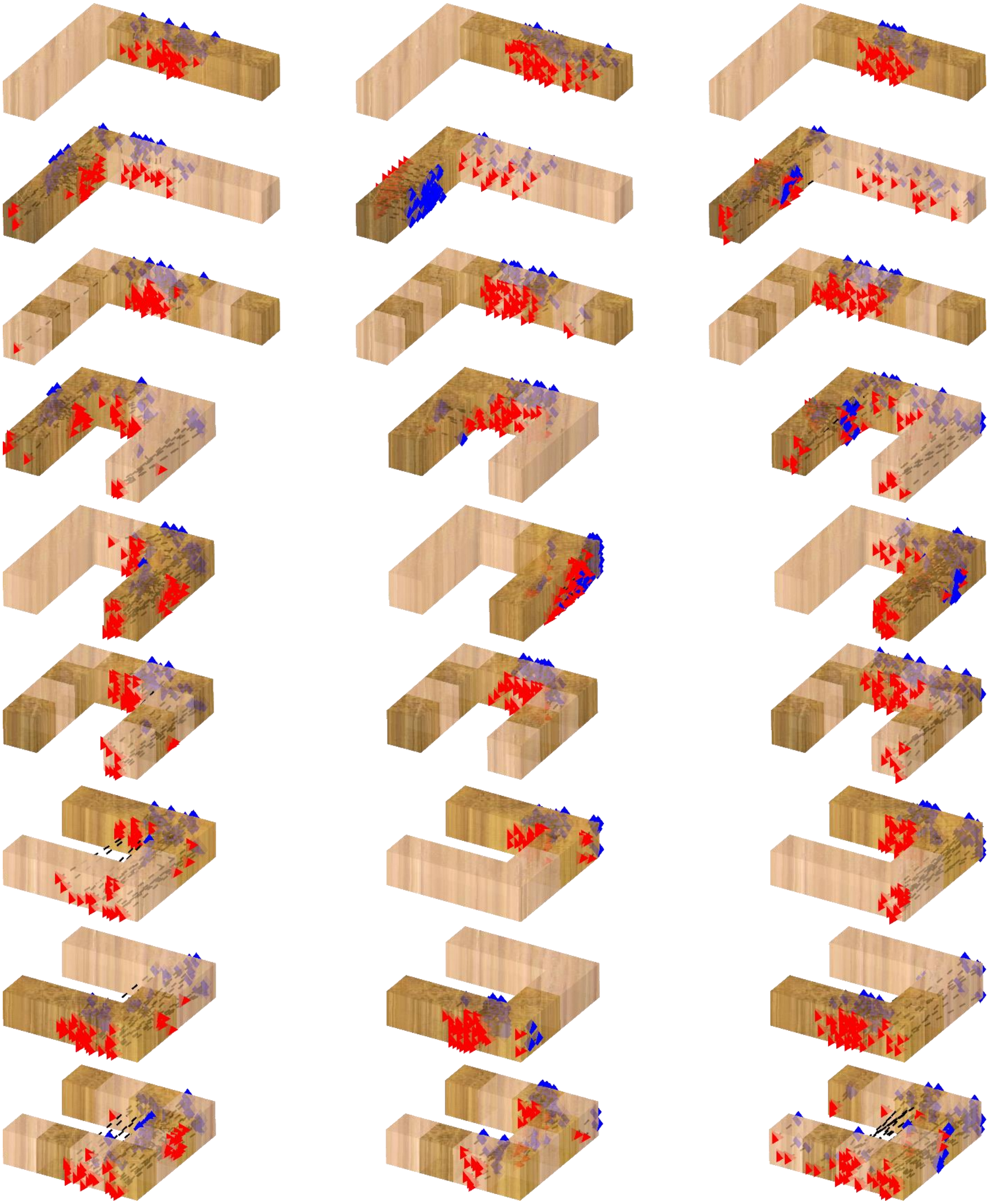

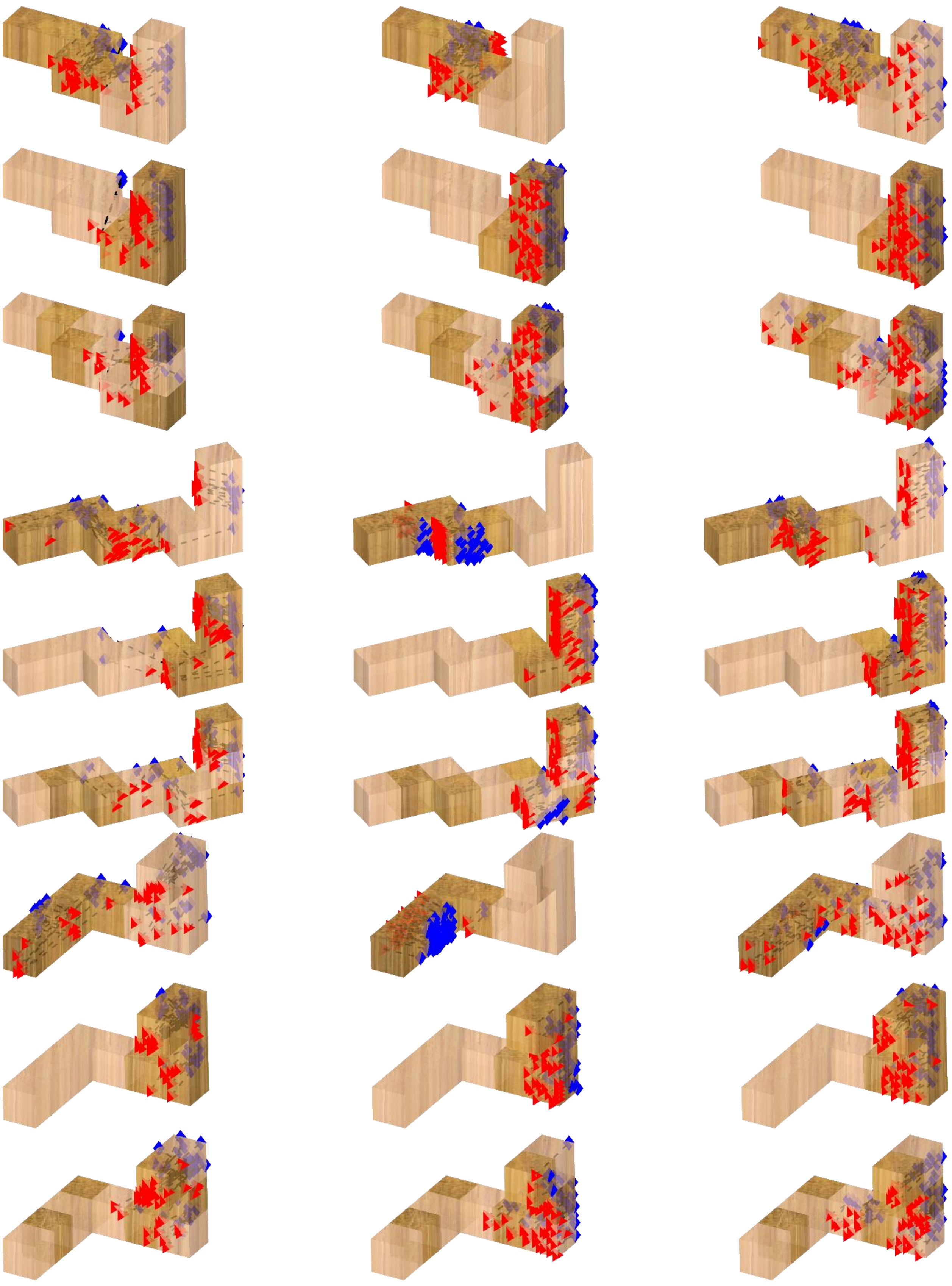

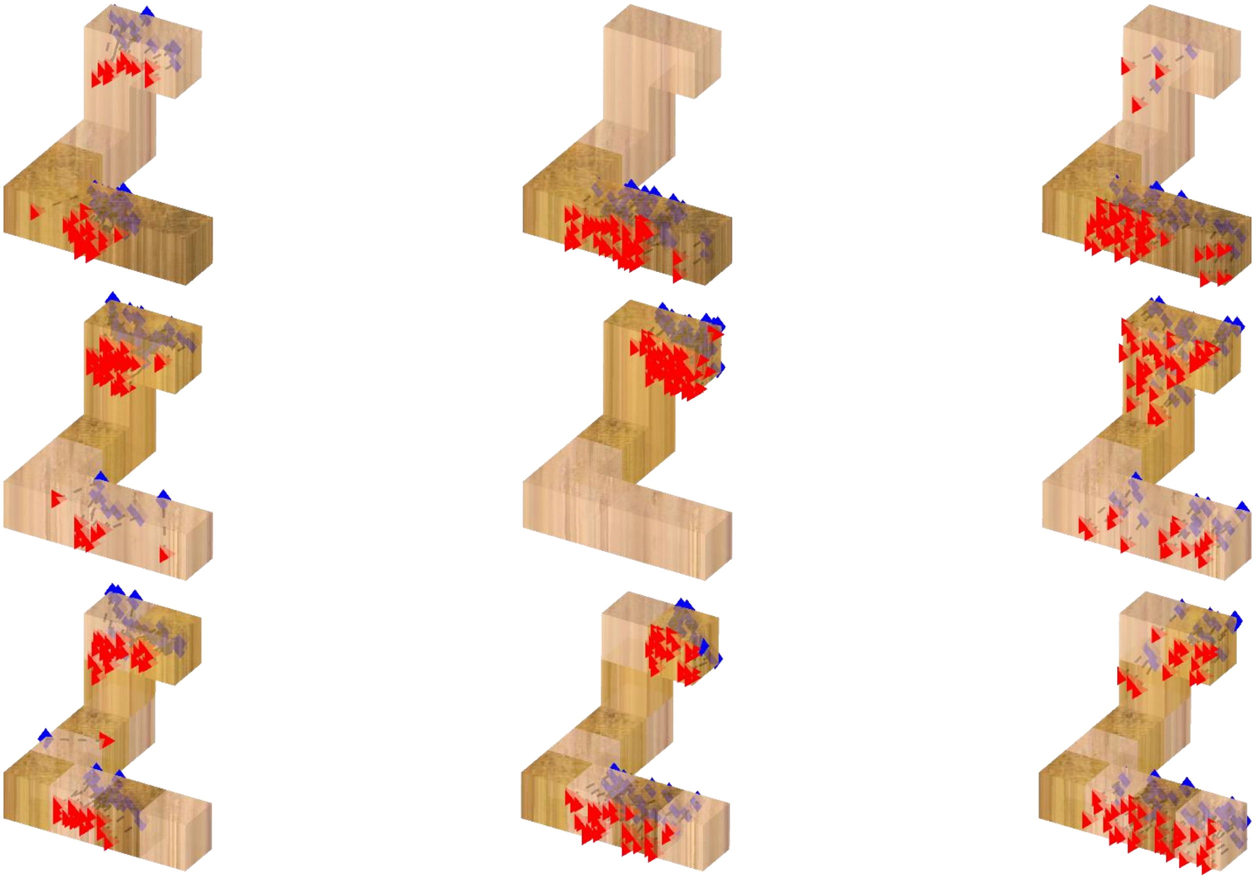

***Figure S1**. Grasping patterns from human participants (left), unfitted model (middle), and fitted model (right). (a) Grasping patterns on wooden objects from Experiment 1. (b) Grasping patterns on mixed material objects from Experiment 2.*
