## Supporting Information S2 Figure for "Predicting precision grip grasp locations on three-dimensional objects"

Department of Experimental Psychology, Justus Liebig University Giessen, Otto-Behaghel-Str.10F, Giessen 35394, Germany


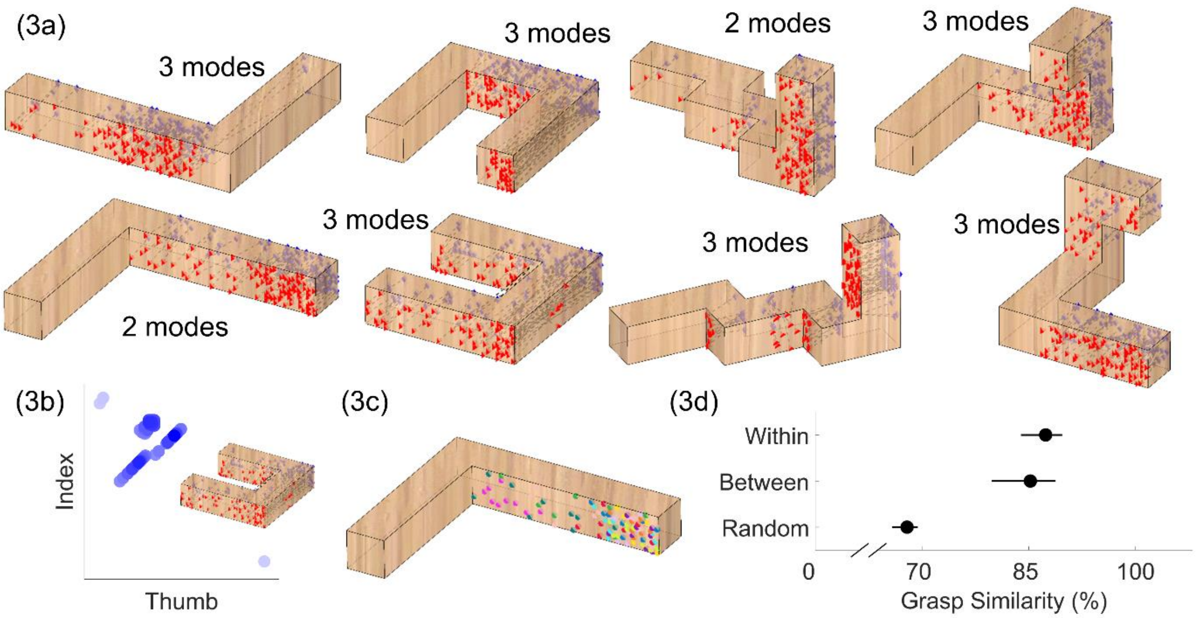

(4a)

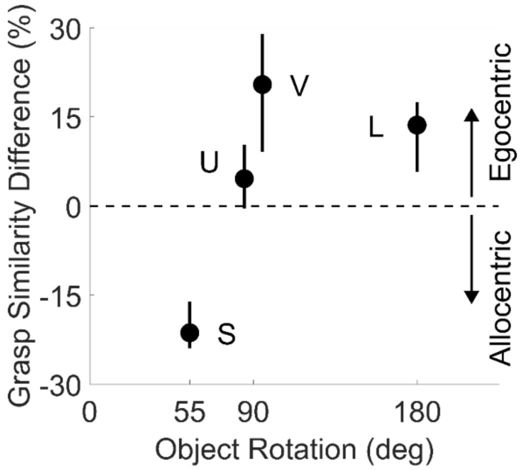

Grasp Similarity Difference (%)
30
15
0
-15
-30
V
U
L
S
Egocentric
Allocentric
0
55
90
180
Object Rotation (deg)


(4b)

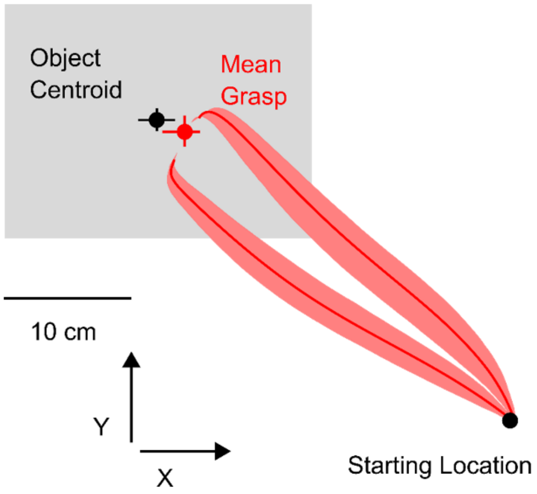

Object
Centroid
Mean
Grasp
10 cm
Y
X
Starting Location


(5a)

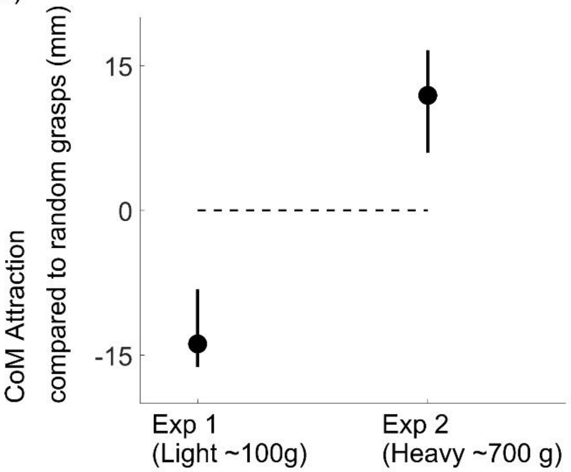

CoM Attraction
compared to random grasps (mm)
15
0
-15
Exp 1
(Light ~100g)
Exp 2
(Heavy ~700 g)


(5c)

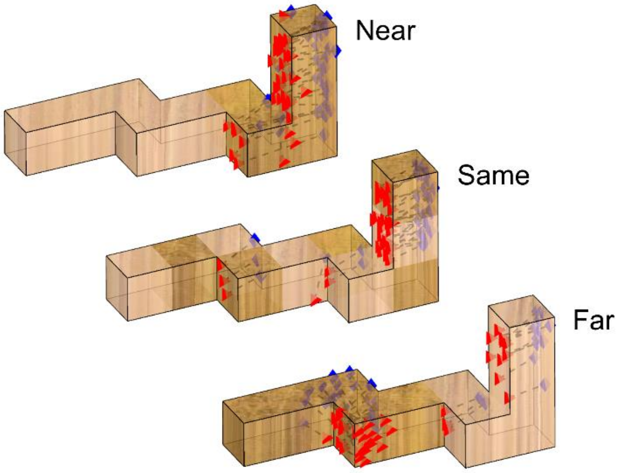

Near
Same
Far


(5d)

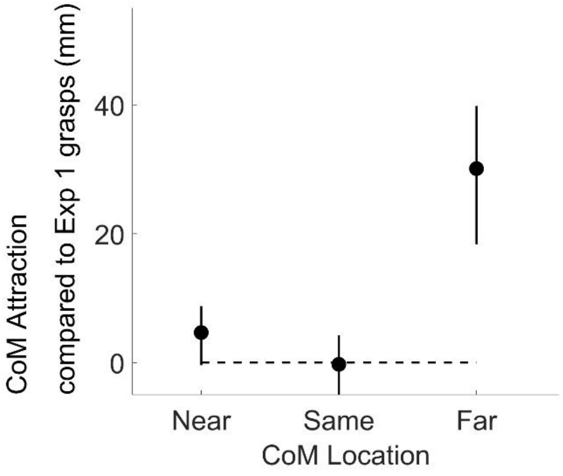

CoM Attraction
compared to Exp 1 grasps (mm)
40
20
0
Near
Same
Far
CoM Location

***S2 Figure. Pattern of empirical results from Experiments 1 and 2 recreated from simulating grasps from the fitted model.*** *Panels are the same as in Figures 3, 4 and 5 of the main manuscript, except that the data are simulated from the model. The grasp trajectories in panel (4b) are from the human data, to highlight how the model correctly reproduces the biases in human grasping patterns. Panel 5b is omitted since the model cannot learn to refine CoM estimates.*
