## Supporting Information S3 Figure for "Predicting precision grip grasp locations on three-dimensional objects"

Department of Experimental Psychology, Justus Liebig University Giessen, Otto-Behaghel-Str.10F, Giessen 35394, Germany

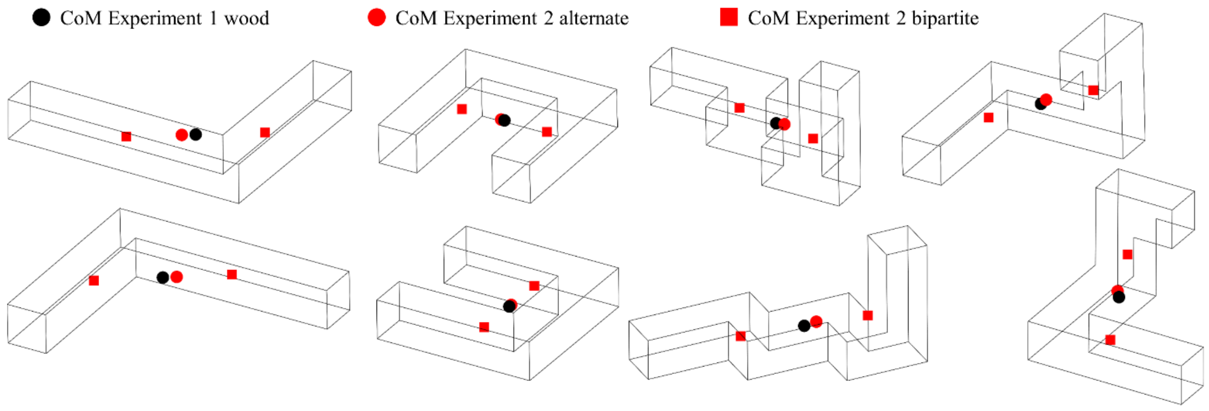


***S3 Figure. Location of the center of mass for the stimuli employed in Experiments 1 and 2.*** *The center of mass of the light wooden objects from Experiment 1 is shown as a black dot. The centers of mass for the heavy alternate and bipartite wood/brass objects from Experiment 2 are shown as red dots and squares respectively.*
